# Mapping multidimensional content representations to neural and behavioral expressions of episodic memory

**DOI:** 10.1101/2023.01.09.523096

**Authors:** Yingying Wang, Hongmi Lee, Brice A. Kuhl

## Abstract

Human neuroimaging studies have shown that the contents of episodic memories are represented in distributed patterns of neural activity. However, these studies have mostly been limited to decoding simple, unidimensional properties of stimuli. Semantic encoding models, in contrast, offer a means for characterizing the rich, multidimensional information that comprises episodic memories. Here, we extensively sampled four human fMRI subjects to build semantic encoding models and then applied these models to reconstruct content from natural scene images as they were viewed and recalled from memory. First, we found that multidimensional semantic information was successfully reconstructed from activity patterns across visual and lateral parietal cortices, both when viewing scenes and when recalling them from memory. Second, whereas visual cortical reconstructions were much more accurate when images were viewed versus recalled from memory, lateral parietal reconstructions were comparably accurate across visual perception and memory. Third, by applying natural language processing methods to verbal recall data, we showed that fMRI-based reconstructions reliably matched subjects’ verbal descriptions of their memories. In fact, reconstructions from ventral temporal cortex more closely matched subjects’ own verbal recall than other subjects’ verbal recall of the same images. Fourth, encoding models reliably transferred across subjects: memories were successfully reconstructed using encoding models trained on data from entirely independent subjects. Together, these findings provide evidence for successful reconstructions of multidimensional and idiosyncratic memory representations and highlight the differential sensitivity of visual cortical and lateral parietal regions to information derived from the external visual environment versus internally-generated memories.

## 1. Introduction

Neuroimaging studies of human episodic memory have found that the contents of memory retrieval are reflected in broadly distributed patterns of neural activity (Danker and Anderson 2010; Rissman and Wagner 2012). While initial fMRI decoding studies of memory focused on relatively coarse information such as the visual category to which a stimulus belongs (Kuhl et al. 2011; Polyn 2005), more recent studies have demonstrated item- or event-specific representations (Favila et al. 2018; Lee et al. 2019; St-Laurent, Abdi, and Buchsbaum 2015; Xiao et al. 2017). However, these studies have overwhelmingly focused on decoding simple, unidimensional, and objective properties of stimuli. In contrast, real-world episodic memories are complex, multidimensional, and subjective (Cooper and Ritchey 2019; Richter et al. 2016). Notably, this limitation is often paralleled in behavioral measures of memory where simple, categorical expressions of retrieval success or accuracy are more common than the kinds of complex and idiosyncratic descriptions humans *actually use* to describe memories (Chen et al. 2017; Gilmore et al. 2021; Heusser, Fitzpatrick, and Manning 2021).

A handful of recent fMRI studies have moved closer toward capturing the richness of memories using multidimensional measures. Naselaris et al. (2015) used an inverted encoding model method (Kay et al. 2008; Naselaris et al. 2011) to reconstruct detailed visual features during mental imagery. Specifically, they mapped low-level visual features extracted from complex natural images to fMRI activity patterns elicited by visual perception. This mapping was then used to successfully predict visual features of independent natural images based on activity patterns evoked during mental imagery. Using a similar approach, Lee and Kuhl (2016) mapped distinct face components to patterns of fMRI activity and then used these mappings to reconstruct faces held in working memory. In another study, Bone, Ahmad, and Buchsbaum (2020) leveraged deep convolutional neural networks to extract visual and semantic features from complex natural images and demonstrated feature-specific reactivation in sensory and frontoparietal cortices during memory retrieval. Collectively, these studies provide important evidence that fine-grained, multidimensional content representations can be mapped to patterns of neural activity evoked during memory retrieval. Notably, however, none of these studies used behavioral measures of memory that matched the richness of the neural measures.

Complementing the studies described above, other fMRI studies have embraced more complex behavioral measures of verbal recall (Chen et al. 2017; Gilmore et al. 2021; Heusser et al. 2021; Nguyen, Vanderwal, and Hasson 2019). For example, Chen et al. (2017) and Nguyen et al. (2019) applied latent semantic analysis (LSA) to verbal recall of movies and Heusser et al. (2021) used topic models to measure changes in verbal recall content over time. Each of these studies found that subject-specific measures of verbal recall content were related to measures of fMRI activity. For example, in Chen et al. (2017) and Nguyen et al. (2019), subjects with more similar recall—or more similar interpretations of the stimuli—showed greater fMRI pattern similarity. In Heusser et al. (2021), the specific time course of content changes during verbal recall was predicted by changes in fMRI activity. While these studies did not directly decode content information from fMRI data, they strongly attest to the feasibility and value of relating subject-specific verbal recall to patterns of neural activity.

To the extent that multidimensional memory representations are captured by patterns of neural activity, an additional question is how these representations are distributed across cortical areas. While memory-based content representations are traditionally viewed as a re-expression of sensory cortical activity (Danker and Anderson 2010), there is now substantial evidence that the lateral parietal cortex (LPC)—a core component of the episodic memory network (Gilmore, Nelson, and McDermott 2015; Rugg and Vilberg 2013)—actively represents the content of retrieved memories (Kuhl and Chun 2014; St-Laurent et al. 2015; Xiao et al. 2017). Moreover, several recent findings specifically suggest that LPC contains the kinds of rich, multidimensional information that is critical for episodic remembering (Bonnici et al. 2016; Cowen, Chun, and Kuhl 2014; Favila et al. 2018; Huth et al. 2016; Lee et al. 2019; Lee and Kuhl 2016; Yu and Shim 2017). There is also emerging evidence for a potential dissociation in content representations across LPC and sensory cortices: whereas content representations in sensory cortex are generally *weaker* during memory retrieval compared to perception, content representations in LPC may be *as strong or stronger* during memory retrieval compared to perception (Favila et al. 2018, 2020; Long and Kuhl 2021; Xiao et al. 2017).

Here, we used semantic encoding models (Kay et al. 2008) and an extensive-sampling fMRI design (thousands of trials per subject) to map multidimensional semantic information from natural scene images to fMRI activity patterns. We then inverted these encoding models (Ester, Sprague, and Serences 2015; Kok, Rait, and Turk-Browne 2020; Sprague, Ester, and Serences 2016) to reconstruct semantic information as subjects viewed and recalled images from memory. These fMRI-based content reconstructions were directly compared to subjects’ verbal recall of the scenes using natural language processing methods. This allowed us to test not only whether fMRI-based reconstructions captured the objective content within scene images, but whether reconstructions matched subjective—and potentially idiosyncratic (subject-specific)— details of how scenes were remembered. Additionally, by comparing reconstructions generated from different regions of visual cortex and LPC, we tested whether these regions differentially expressed content information during image viewing versus image recall. Finally, we tested whether semantic encoding models successfully generalized across subjects—a question that has important implications for leveraging data-rich models from extensively-sampled subjects.

## 2. Materials and Methods

### 2.1. Subjects

Nineteen experimental sessions were collected from four human subjects (two females, age 23-30 years) from the University of Oregon community. Three subjects completed five sessions each and one subject completed four sessions. The sample size was modeled after Naselaris et al. (2015), which used a similar encoding model procedure for memory-based reconstructions. Despite the small sample size, each subject was sampled extensively across a large number of stimuli, a procedure which may have distinct advantages compared to sampling many individuals across a more limited number of stimuli (Naselaris, Allen, and Kay 2021). All subjects were right-handed and reported normal or corrected-to-normal vision. Informed consent was obtained in accordance with procedures approved by the University of Oregon Institutional Review Board.

### 2.2. Stimuli

Two sets of image stimuli were prepared: one for use in a recognition memory task and one for use in a recall memory task. The recognition set contained a total of 5000 complex scene images, which were selected from the Microsoft COCO dataset (http://cocodataset.org/, Lin et al., 2015). These images depict complex everyday scenes of common objects from 91 categories in their natural context. Each image in the dataset is annotated with five written descriptions from independent human subjects. These descriptions capture the main content of the images and were used, in the present study, as information channels for the inverted encoding model. For each subject and each session, 680 images were randomly selected (without replacement) from the recognition set. Of these, 600 were studied prior to the fMRI session and served as ‘old’ items in the recognition test. The remaining 80 images served as novel foils (‘new’ items) in the recognition test. The recall set consisted of a total of 100 images, also drawn from the Microsoft COCO dataset. Each image was randomly paired with a texture (taken from the internet), creating a set of paired associates. The textures served as cues during the cued recall task (described below). For each session, 20 of the paired associates were studied and tested. The same 20 pairs were used in each session for each subject in order to facilitate across-subject analyses.

### 2.3. Experimental design and procedures

#### Overview of paradigm

Each session of the experiment consisted of two separate parts which were conducted across consecutive days (**Fig. 1A**). On Day 1 of each session, subjects were overtrained on 20 paired associates (textures + scene images) and were familiarized with a separate 600 scene images (from the recognition set). On Day 2 of each session, subjects first completed additional training on the paired associates from Day 1 and additional familiarization with images from the recognition set. Then, during fMRI scanning subjects completed two phases: (1) a covert cued recall phase in which the 20 textures were repeatedly used to test memory for the associated scenes, and (2) a recognition memory phase which included the 600 familiarized images + 80 lures. Finally, subjects exited the scanner and completed an overt (verbal) cued recall test for the 20 paired associates. This two-day procedure constituted a single session and each subject completed 4-5 sessions. In order to minimize across-session memory interference, there was a delay of at least 7 days between sessions, for each subject.

**Fig. 1.**
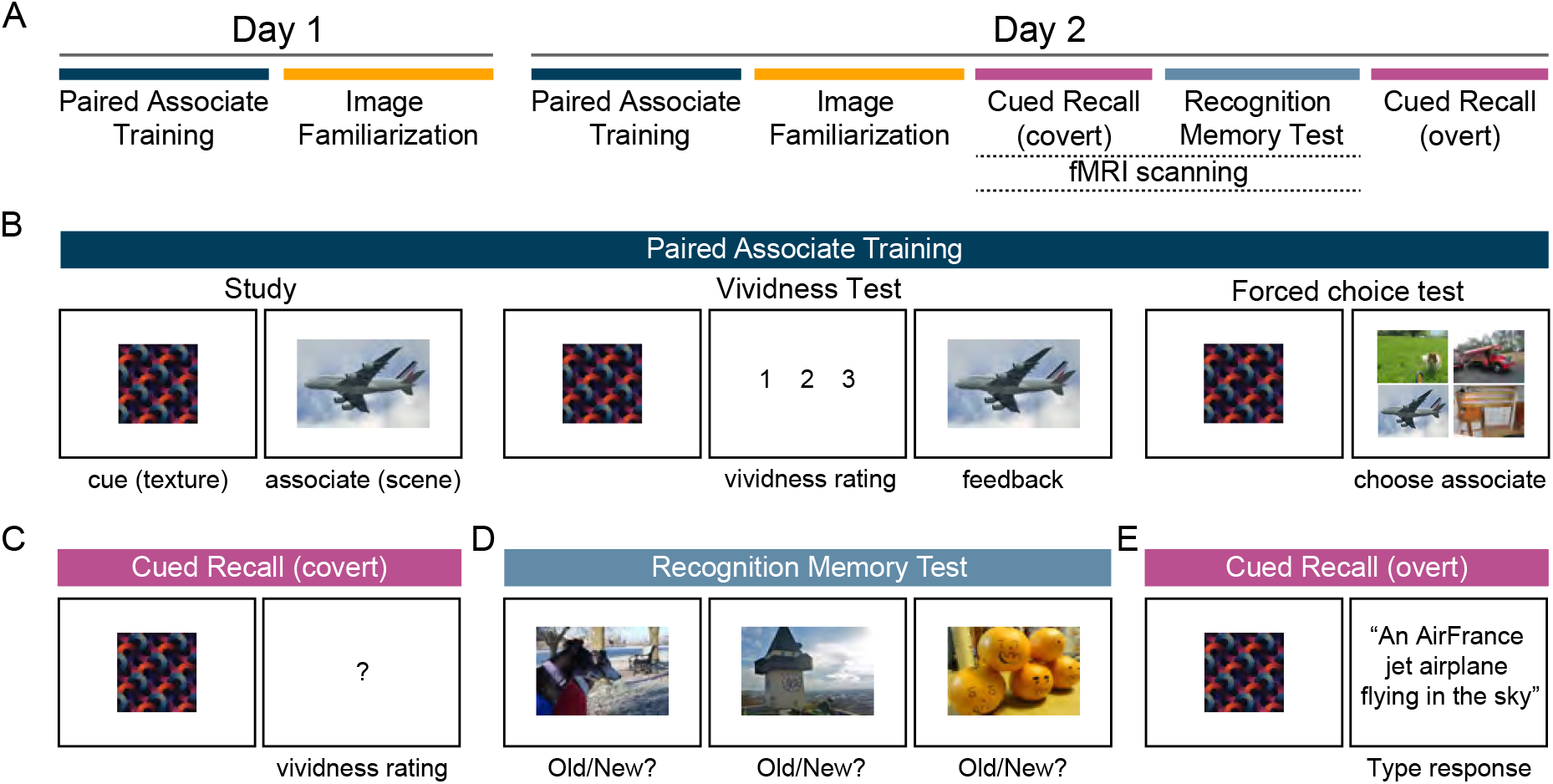
Experimental procedure. **A**. Overview of experimental phases. Each subject completed 4 to 5 experimental sessions. Each experimental session involved two consecutive days of tasks. On Day 1, subjects learned 20 associations between cues (textures) and associates (scenes) via a paired associate training procedure and were also familiarized with 600 additional scene images (image familiarization). No fMRI scanning was conducted on Day 1. On Day 2, subjects completed additional paired associate training and image familiarization before entering the scanner. During scanning, subjects completed a covert cued-recall test of the cue-associate pairs followed by a recognition memory test. After exiting the scanner, subjects completed an overt cued-recall test for the cue-associate pairs. **B**. The paired associate training included three phases: study, vividness test, and forced choice test. During study, subjects were shown textures followed by scenes and attempted to learn these associations. During the vividness test, subjects were shown textures and then indicated the vividness with which they were able to recall the corresponding scene. The correct scene was then shown as feedback. During the forced choice test, subjects were shown a texture followed by four previously-studied scenes and were asked to select the corresponding scene. **C**. In the scanned (covert) cued-recall phase, subjects were shown textures and rated the vividness with which they could recall the corresponding scene (as in the vividness test, but without feedback). **D**. In the scanned recognition memory test, subjects made old/new judgements for scenes (that did not include the scenes from the paired associate training). The sole purpose of the recognition memory test was to train the semantic encoding models. **E**. In the final (overt) recall test, subjects were shown textures and typed a sentence to describe the content of the corresponding scene.

#### Paired associate training

Paired associate training was conducted at the beginning of Day 1 (4-5 rounds) and the beginning of Day 2 (1 round). Each round of paired associate training consisted of three distinct phases in the following sequence: study, vividness test, study, vividness test, forced choice associative memory test. In the study phases, subjects saw and deliberately encoded each of the 20 paired associates, one pair at a time. On each trial, the cue (texture) was first presented for 1 s, followed by a fixation cross for 0.5 s, and then the target image (scene) for 2 s. Another fixation cross was presented for 1 s at the end of each trial (before the start of the next trial). In the vividness test phases, each cue was presented for 1 s followed by a 3-point vividness scale (“1 2 3”) and subjects reported, via button press, the vividness with which they could recall the target image (1-*Can’t remember*, 2-*Remember*, and 3-*Vividly remember*). The vividness report was self-paced. After responding, feedback was given by presenting the target image alone on the screen for 1.5 s. A fixation cross was presented for 0.5 s in between trials. In the forced choice associative memory test, a cue image was first presented for 1 s and then, after a fixation cross (0.5 s), four scene images appeared on the screen. The four images included the target (correct) scene along with three scenes randomly selected from the remaining 19 scenes in the set of paired associates studied in the current session. Subjects were instructed to select the scene image that had been paired with the cue by pressing one of four keys. There was no time limit to respond. After subjects made a selection, feedback was provided. If the correct image was selected, a green fixation cross was presented (0.5 s) followed by the correct image presented in the center of the screen (1 s). If an incorrect image was selected, a red fixation cross was presented (0.5 s) followed by the correct image (1 s) presented in the center of the screen. Finally, a black fixation cross was presented for 1 s (until the start of the next trial). For each session, subjects were required to reach at least 95% accuracy for two consecutive rounds on Day 1 before proceeding to the Image Familiarization Phase. Using this performance criterion, all subjects completed 5 paired associate training rounds on Day 1 for each session, with the exception of one subject that reached the criterion by the 4^th^ round for one of the sessions.

#### Image Familiarization

For each session, Image Familiarization was conducted on Day 1 and Day 2, immediately after the paired associate training rounds. During each familiarization phase subjects saw all 600 scene images presented in the center of the screen, one at a time and in random order, and distributed across five blocks (120 images/block). Subjects were instructed to try their best to remember each image for a later memory test (the recognition memory test). No behavioral responses were made. On Day 1, each image was presented for 1 s with a 0.5 s fixation cross in between trials. On Day 2, each image was presented for 0.6 s with a 0.4 s fixation point in between trials.

#### Scanned Cued Recall

For each session, subjects completed two fMRI runs of a covert cued recall task (**Fig. 1C**), each lasting 6 min and 16 s. As during the paired associate training rounds, subjects were shown cues (textures) and indicated the vividness with which they could recall the corresponding scene image using a 3-point scale. Each run consisted of 40 recall trials (2 trials per association per run), with the order of trials in each run pseudorandomized with the constraint that the same association was not tested consecutively. Every trial started with a cue image centrally presented over a white background for 0.5 s. Next, a question mark appeared in the center of the screen (3.5 s), prompting subjects to make their vividness response using a button box. Finally, a fixation cross was presented either for 4 s (75% of trials) or 8 s (25% of trials).

#### Scanned Recognition Memory Test

Following the cued recall task, subjects completed the recognition memory test (**Fig. 1D**) which consisted of eight runs, each lasting 6 min and 20 s. Each run contained 75 old images and 10 novel images presented in random order, for a total of 680 images across the 8 runs. Each trial began with the presentation of a scene in the center of the screen (1 s). Next, a question mark (3 s) prompted subjects to make an old/new decision by pressing one of two keys on a button box. After a small number of the recognition trials (6/85), a fixation cross was presented for 4 s. The rationale for including a disproportionate number of old images (600 out of 680) was because fMRI data from the recognition memory test was used to train encoding models applied to the cued recall task and we sought to increase the extent to which these models were trained on ‘memory data’ (old trials). Specifically, recent evidence indicates systematic differences in the spatial activity patterns associated with memory-based content representations compared to perception-based content representations (Favila et al. 2020). Thus, our intuition—though not a point we directly tested—was that transfer from the recognition to recall trials might benefit from the recognition trials having a high percentage of old trials. Additionally, the recognition memory test served as a cover task to help keep subjects engaged while viewing hundreds of images per session.

#### Post-scan Cued Recall

After subjects exited the scanner, they completed a final cued recall test (**Fig. 1E**). However, in contrast to the prior cued recall tests which recorded covert (vividness) memory judgments, here subjects were asked to explicitly describe their memories. On each trial, subjects were shown a cue (texture) and were asked to type a sentence to describe the content of the associated scene image. Specifically, the instructions asked subjects to “write a complete but simple sentence” that should “include adjectives if possible, describe the main characters, the setting, or the relation of the objects in the image, and try to be concise”. After subjects typed their response on the computer screen, they pressed enter to advance to the next trial. No time limit was given and each of the 20 associations from the current session was tested once, in random order.

### 2.4. fMRI data acquisition

fMRI scanning was conducted on a Siemens 3 T Skyra scanner at the Robert and Beverly Lewis Center for NeuroImaging at the University of Oregon. Before the functional imaging, a whole-brain high-resolution anatomical image was collected for each subject and each session using a T1-weighted protocol (grid size 256 × 256; 176 sagittal slices; voxel size 1 × 1 × 1 mm). Whole-brain functional images were collected using a T2*-weighted multi-band accelerated EPI sequence (TR = 2s; TE = 25ms; flip angle = 90°; 72 horizontal slices; grid size 104 × 104; voxel size 2 × 2 × 2 mm). Each cued recall scan consisted of 188 volumes. Each recognition memory test scan consisted of 190 volumes.

### 2.5. fMRI data preprocessing

MRI data were first converted to Brain Imaging Data Structure (BIDS) format using in-house scripts. MRIQC v0.15.1 (Esteban et al. 2017) was used for preliminary data quality assessment. We applied a threshold that no more than 20% of TRs in any scan run could exceed a framewise displacement of 0.3 mm; however, no scan runs were excluded using this threshold. Preprocessing was performed using FMRIPrep v1.5.4 (RRID:SCR_016216) (Esteban et al., 2019), a Nipype (RRID:SCR_002502) based tool, with the default processing steps. Each structural image was corrected for intensity non-uniformity and skull-stripped. Brain surfaces were reconstructed using recon-all from FreeSurfer v6.0.1. Spatial normalization to the ICBM 152 Nonlinear Asymmetrical template version 2009c was performed through nonlinear registration with the antsRegistration tool of ANTs v2.1.0, using brain-extracted versions of both T1w volume and template. Brain tissue segmentation of cerebrospinal fluid (CSF), white-matter (WM) and gray-matter (GM) was performed on the brain-extracted T1w using FAST (FSL v5.0.9).

Functional data were slice time corrected, motion corrected, and corrected for field distortion. This was followed by co-registration to the corresponding T1w using boundary-based registration with six degrees of freedom using bbregister (FreeSurfer v6.0.1). Motion correcting transformations, BOLD-to-T1w transformation and T1w-to-template (MNI) warp were concatenated and applied in a single step using antsApplyTransforms (ANTs v2.1.0) using Lanczos interpolation. We then applied a high pass filter using a cutoff period of 100 s. Finally, the preprocessed fMRI data were smoothed by a 1.6 mm full-width-half-maximum Gaussian kernel with FSL’s SUSAN (Smoothing over Univalue Segment Assimilating Nucleus) (Smith and Brady 1997). Grand-mean intensity normalization of each functional image volume was performed by a single multiplicative factor. Confounding regressors including framewise displacement (FD), global signal, white matter, and cerebrospinal fluid signals were generated for each volume. Within-subject reconstructions were conducted in subjects’ native EPI space, and across-subject reconstructions were conducted in the standard space.

### 2.6. Regions of interest

Regions of interest (ROIs) included four subregions of the posterior lateral parietal cortex (LPC), the ventral temporal cortex (VTC), and the occipital temporal cortex (OTC) (**Fig. 4A**). ROIs were defined using FreeSurfer’s Destrieux atlas (the following label numbers refer to Simple_surface_labels2009.txt). The subregions of LPC consisted of the angular gyrus (ANG, #25), supramarginal gyrus (SMG, #26), superior parietal lobule (SPL, #27), and intraparietal sulcus (IPS, #57). The VTC ROI was comprised of regions 21, 23, 51, 52, 61, and 62. The OTC ROI was comprised of regions 2, 19, 43, 58, and 60. The ROIs were co-registered to the functional images and further masked by subject-specific whole-brain masks generated from functional images to exclude areas where signal dropout occurred. All ROIs contained brain regions from both hemispheres (mean number of voxels for each ROI: 1816 in ANG; 1942 in SMG; 1594 in SPL; 1318 in IPS; 4459 in VTC; 3854 in OTC).

### 2.7. Single-item response estimation

For each session, separate general linear models (GLM) were created for each of the 20 images during the cued recall task and each of the 680 images during the recognition memory test. A least-square single method was used for each item, where the given item was modeled with a single regressor and all the remaining items were modeled with another regressor. The presentation of each stimulus was modeled as an impulse and convolved with a canonical hemodynamic response function (double gamma). The GLM included head-motion parameters (six rotation and translation head movement estimates) and nuisance regressors marking outlier TRs (FD > 0.3 mm from previous TR) as confounding regressors. The *t*-statistic values associated with each image were used in the semantic encoding model to increase reliability by noise normalization (Walther et al. 2016).

### 2.8. Image content reconstruction

To represent the content of each scene image, we used the Word2vec embedding algorithm. This algorithm transforms single words into 300-dimensional vectors (word embeddings). Similarities/distances between these vectors reflect the similarity of the corresponding words. In our analysis, we took advantage of the annotation captions (five captions for each image) from the COCO dataset. After a standard preprocessing procedure that included filtering of stop words and tokenization, we obtained the word embeddings for the critical words in the annotation captions. We calculated the mean vector, across the five captions, to represent the content in each image (**Fig. 2A**). We then applied principal component analysis (PCA) on the entire pool of 300-dimensional word embeddings for the 5100 images (i.e., the full set of recognition + recall images). The first 30 components, which explained 68.59% of the total variance were used as information channels in the semantic encoding model. We refer to these 30 components as *semantic component scores*. The goal of reconstruction analyses was to accurately predict the semantic component scores.

**Fig. 2.**
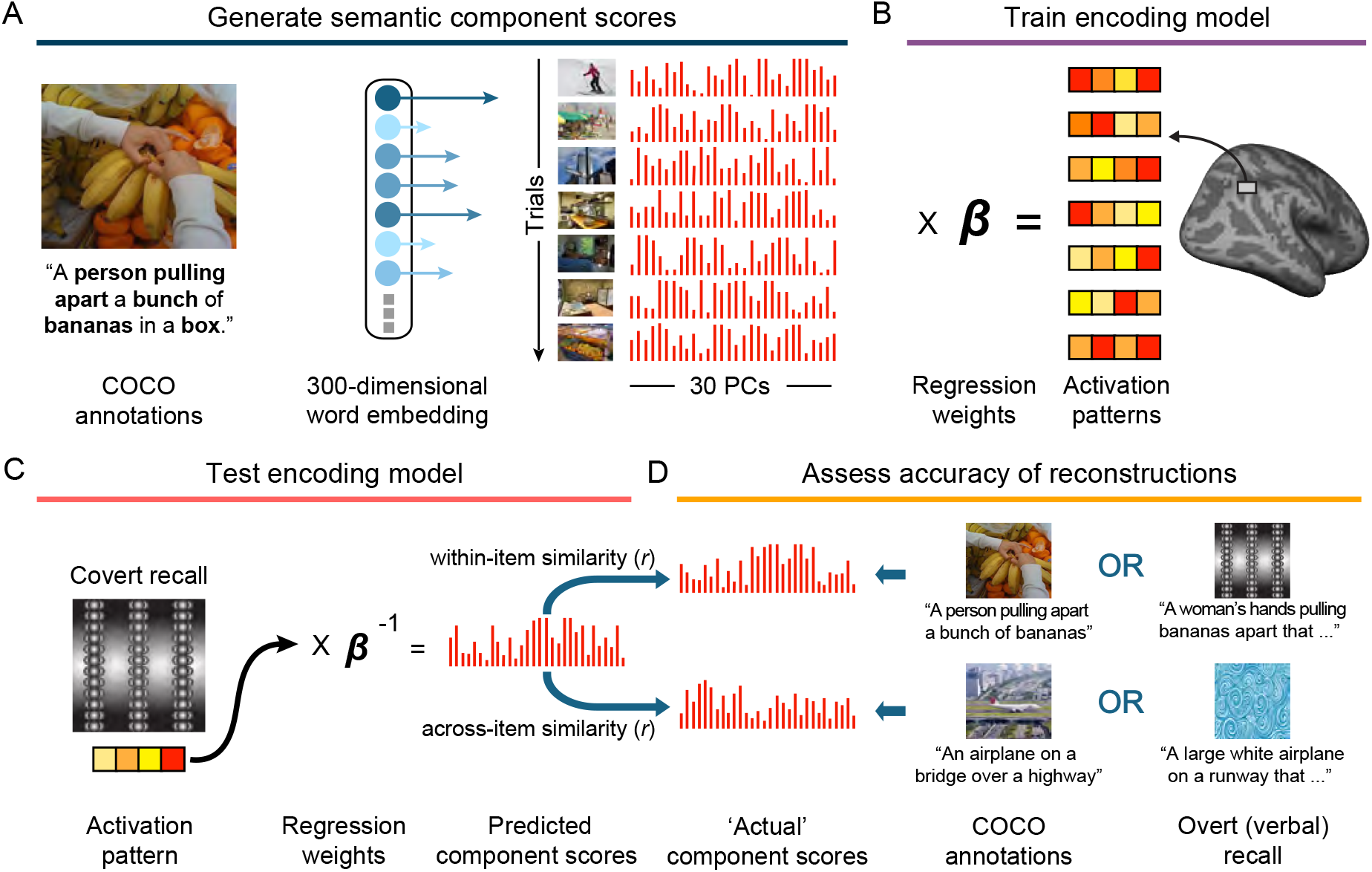
Schematic overview of the semantic content reconstruction analysis. **A**. Generating semantic component scores. Annotations from the COCO image dataset were used as semantic descriptions of the images. After filtering out the stop words, the captions were transformed into 300-dimensional vectors using the Word2Vec embedding method. PCA was run on all of the 5100 candidate images, and the first 30 principle components (hereinafter, semantic components) were extracted so that the content of each image could be expressed as a weighted sum of these components. **B**. Training the encoding model. Linear regression was used to estimate a model that learned the mapping between the semantic component scores of the trained images (i.e., the training set) and the fMRI activation patterns they evoked. **C**. Testing the encoding model. The regression weights obtained from the training set were applied to an independent set of images (i.e., the testing set) to predict semantic component scores. Encoding models were tested using cued recall trials (shown) or recognition trials (not shown). **D**. Assessing reconstruction accuracy. The accuracy of reconstruction for each image was determined by computing the Pearson correlations between the predicted semantic component scores and the actual semantic component scores. Actual semantic component scores were either based on the COCO dataset captions (left side of boxes) or the verbal recall responses subjects generated in the final cued-recall test (right side of boxes). Correlations were separately computed for ‘matching’ images (within-item similarity) and non-matching images (across-item similarity). Reconstructions were considered accurate if within-item similarity was higher than across-item similarity.

Reconstructions of semantic component scores were generated using a cross-validation approach. fMRI activation patterns evoked during the recognition trials were used as training patterns to estimate the relationship between fMRI activity patterns and semantic component scores (**Fig. 2B**). We modeled the response in each voxel as a weighted sum of the information channels (i.e., the 30 semantic components):

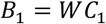

where *B*_1_ (n images × m voxels) is the activation patterns of voxels (*t* maps) during the recognition memory test, *C*_1_ (n images × k components) is the modeled response of each component, or information channel, on each trained image, and *W* (k components × m voxels) is a weight matrix quantifying the contribution of each information channel to each voxel (**Fig. 2B**). We can solve for *W* using the following ordinary least-squares linear regression:

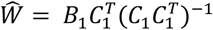

Given the estimated weights within an ROI (*Ŵ*) and a novel pattern of activation (*B*_2_) from the recognition trials (recognition-based reconstruction) or the recall trials (recall-based reconstruction), we can compute an estimate of the semantic component scores by inverting the model (**Fig. 2C**):

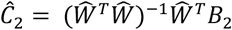

#### Recognition-based reconstruction

Separately for each subject, N-fold cross-validation was performed on recognition data where N equals the number of scanning runs pooled across all of the sessions that each subject completed (i.e., 40 runs for the three subjects that completed 5 sessions each and 32 runs for the remaining subject that completed 4 sessions). For each fold, the activation patterns from N-1 runs (i.e., 39 or 31 runs) were used as training patterns and those of the remaining run served as the testing set (i.e., the trials for which the semantic component scores were predicted). In this manner, all trials iteratively contributed to both model training and model testing.

#### Recall-based reconstruction

To predict semantic component scores during recall trials, activation patterns from recognition runs were used as training data and the estimated weights based on the recognition trials (training data) were then applied to the recall trials (testing data) to predict the semantic component scores for each of the recalled images. Recall-based reconstructions were performed in two ways: within-subjects and across-subjects. For within-subject reconstructions, all of the recognition runs across all sessions for a given subject were used as the training data and the testing data were all of the recall runs across all sessions for that same subject. For across-subject reconstructions, all of the recognition runs across all sessions from N-1 subjects were used as the training data and the testing data were all of the recall runs across all sessions from the held-out subject.

### 2.9. Reconstruction accuracy

For all reconstruction analyses, accuracy was based on Fisher-transformed Pearson correlations between predicted (reconstructed) and ‘actual’ semantic component scores. ‘Actual’ semantic component scores were either based on COCO annotations (which were derived from an independent set of subjects) or from subjects’ verbal recall responses (which were collected in the final cued recall test, after fMRI scanning). Unless otherwise noted, successful reconstruction accuracy was defined as greater *within-item correlations* than *across-item correlations* (**Fig. 2D**). Within-item correlations refer to correlations between reconstructed and actual semantic component scores corresponding to the same image. Across-item correlations refer to the mean of correlations between reconstructed and actual semantic component scores corresponding to different images [e.g., *r*(reconstructed scores for image 1, actual scores for image 2)]. Across-item correlations were always restricted to images from the same fMRI session. Additionally, within-item correlations for recognition-based reconstructions were only compared against across-item correlations for other recognition-based reconstructions; likewise, within-item correlations for recall-based reconstructions were only compared against across-item correlations for other recall-based reconstructions. Group-level results were obtained by first averaging correlations within sessions for each subject and then across sessions and subjects.

### 2.10. Statistical analysis

Unless otherwise stated, mixed-effects models were used to test the reconstruction accuracy of correlation difference measures. Linear mixed-effects models were implemented with lme4 in R 3.6.3, fitted using restricted maximum likelihood. To determine whether within-item correlations differed from across-item correlations, we used the likelihood ratio test to compare models with (full model) and without (null model) the predictor of interest (i.e., correlation type: within-item correlation or across-item correlation). Subject and session numbers were included as random factors. For statistical tests of reconstruction accuracy within individual ROIs, uncorrected *p* values are reported. In tests that compared reconstruction accuracies across ROIs or conditions, mixed-effects models were used with the subject number and session number included as random factors.

## 3. Results

### 3.1. Behavioral performance

Group-level results were obtained by first averaging data within sessions for each subject and then across sessions and subjects. On Day 1 of each session, subjects studied 20 paired associates (textures with scenes) across 4-5 rounds. For each round, memory was tested via forced-choice associative memory and cued recall tasks. In the forced-choice associative memory test, subjects were asked to select the target image for each cue from a set of three image choices. Performance was high across all rounds (Round 1: 97.89% ± 4.19%; Round 2: 98.95% ± 2.09%; Round 3: 98.68% ± 2.27%; Round 4: 98.68% ± 2.81%; Round 5: 97.81% ± 3.15%; **Fig. 3A**). In the cued recall tasks, subjects reported how vividly they were able to recall the target image on a 3-point scale. The mean percentage of “Vividly remember” responses (the highest rating) was 49.74% ± 22.96% (SD) in round 1, 93.16% ± 10.23% in round 2, 99.34% ± 1.83% in round 3, 99.74% ± 0.79% in round 4, and 100% ± 0.00% in round 5 (mean rescaled responses are shown in **Fig. 3B**). Critically, performance remained high for both tasks at Day 2 as evidenced by performance on the forced-choice associate memory test that occurred just prior to scanning (Round 6: 98.95% ± 2.09%; **Fig. 3A**) and the rate of “Vividly remember” responses during the cued recall tasks that occurred just prior to scanning (Round 6: 98.16% ± 3.89%) and during scanning (scan: 98.55% ± 6.00%) (mean rescaled responses are shown in **Fig. 3B**)

**Fig. 3.**
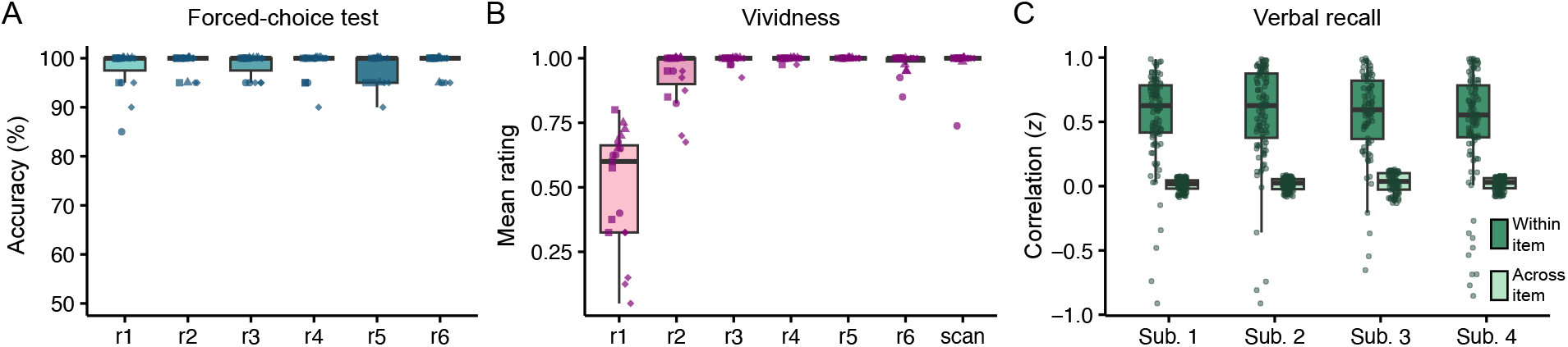
Behavioral performance across the entire experimental procedure. **A**. Forced-choice test accuracy was measured during the paired associate training rounds on Days 1 and 2. The first five rounds (r1–r5) were completed during Day 1. The 6^th^ round (r6) was completed during Day 2 (just prior to fMRI scanning). Chance accuracy = 25%. **B**. Vividness ratings were made during the first five paired associate training rounds on Day 1 (r1–r5), during the 6^th^ paired associate training round on Day 2 (r6), and during the covert cued recall test during fMRI scanning on Day 2 (scan). Ratings were rescaled from 1, 2, 3 to 0, 0.5, 1.0 with 0 corresponding to the lowest vividness rating and 1.0 to the highest vividness rating. For **A** and **B**, data are represented by boxplots with dots representing data from individual sessions with each subject represented by a different shape. Note: for many of the rounds performance was at ceiling and boxplots are therefore compressed. **C**. Verbal recall performance from the overt cued recall test following scanning on Day 2. For each subject (Sub. 1–4) and each recalled image, Pearson correlations were computed between the 30 semantic components generated from subjects’ verbal responses and semantic components generated from the independent COCO annotations of (i) the same images (within-item similarity) or (ii) other images from the recall set (across-item similarity). For within-item similarity, each dot represents the within-item correlation for a single recall trial to its corresponding COCO annotations. For across-item similarity, each dot represents the mean *z*-transformed correlation between a single recall trial and all non-corresponding COCO annotations.

**Fig. 4.**
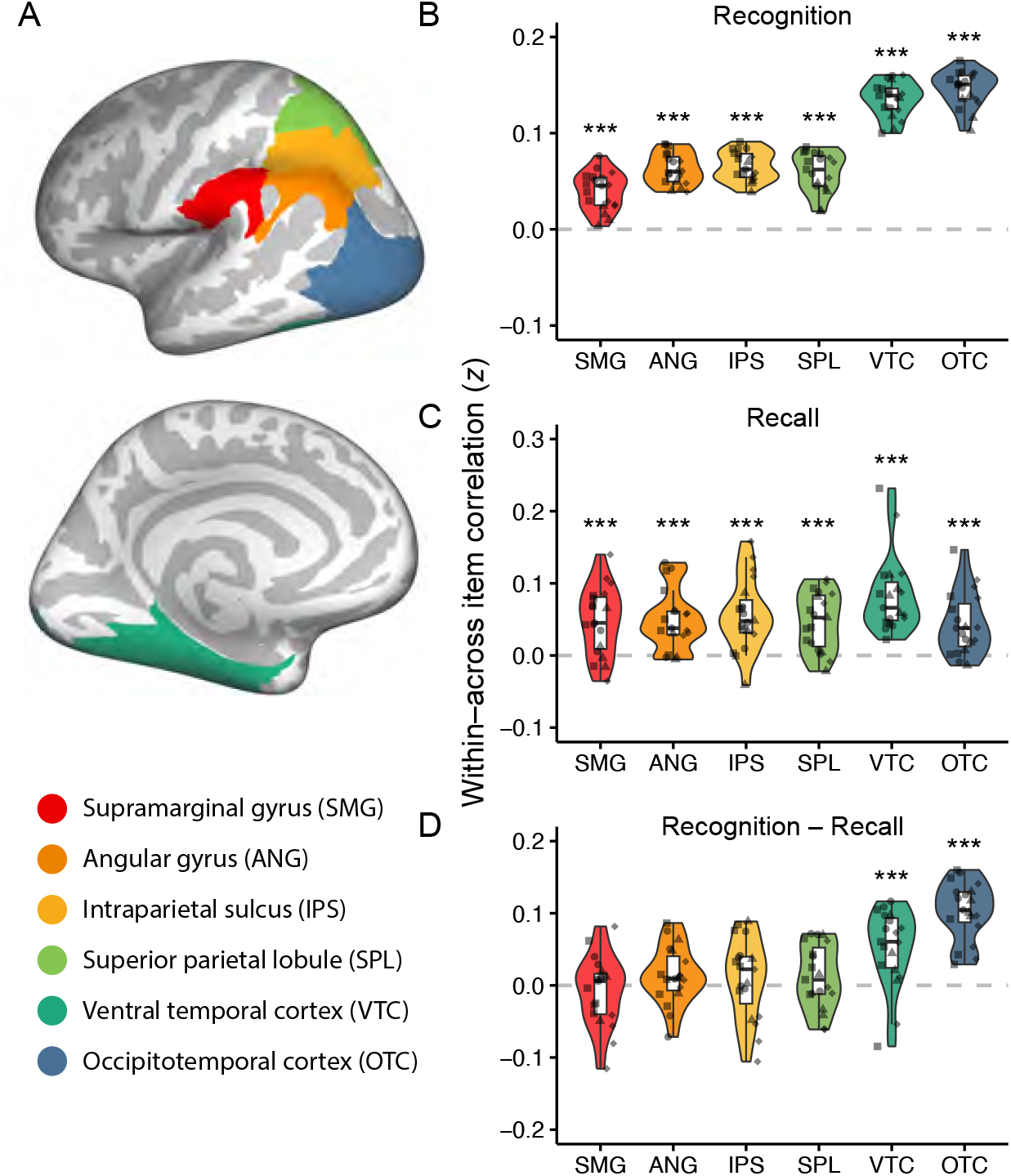
Accuracy for fMRI-based reconstructions of semantic component scores. **A**. Anatomical regions of interest (ROIs), visualized on the inflated surface of an averaged template brain (from FreeSurfer). Top: left lateral view. Bottom: left medial view. **B**. Mean reconstruction accuracies of semantic component scores for each ROI based on encoding models trained and tested on recognition trials. Independent COCO annotations were used to define the ‘actual’ content of each image and semantic component scores from these annotations were then compared to semantic component scores reconstructed from fMRI activity patterns during the covert cued recall phase. Accuracy is expressed as within-item correlations – across-item correlations, with positive values (i.e., > 0) reflecting successful (item-specific) reconstructions. Accuracy was significantly above chance for all ROIs. **C**. As in **B**, but based on encoding models trained on recognition trials and tested on recall trials. Accuracy was significantly above chance for all ROIs. **D**. Difference in reconstruction accuracy for recognition vs. recall trials (**B** vs. **C**). Positive values reflect higher accuracy for recognition trials than recall trials. Only VTC an OTC exhibited significantly greater accuracy for recognition-based reconstructions than recall-based reconstructions. Notes: dots represent data from individual sessions with each subject represented by a different shape; *** *p* < 0.001.

After exiting the scanner, subjects completed a final post-scan cued recall task during which they generated a sentence to describe the content in each target image. These subject-specific recall-based descriptions were transformed to 300 dimensional vectors (word embeddings) using Word2Vec. The COCO annotations for each of these images were also transformed to word embeddings using Word2Vec. We then calculated the Pearson correlations between the word embeddings from subjects’ verbal recall and those from the independent COCO annotations. As shown in **Fig. 3C**, each subject exhibited markedly higher within-item correlations (i.e., correlations between verbal recall vectors and COCO annotation vectors corresponding to the same image) than across-item correlations (i.e., correlations between recall vectors and annotation vectors corresponding to different images). These results confirm that subjects were able to accurately describe images from memory and also validate our approach of characterizing verbal recall through word embeddings.

For the recognition memory test conducted during scanning, mean recognition sensitivity (d’) across sessions and subjects was 1.98 ± 0.52. The mean hit rate for studied images was 84.63% ± 11.51%, and the mean correct rejection rate for new images was 76.97% ± 12.48%. A mixed effects model including subject and session numbers as random factors showed that the hit rate was significantly higher than the false alarm rate (χ^2^_1_ = 89.96, *p* < 0.0001, β = 0.616, SE = 0.030).

### 3.2. Reconstruction of content from viewed images

For fMRI analyses, we first tested for successful reconstruction of image content from activity patterns evoked in visual and lateral parietal cortices during the recognition memory task (when images were visually presented on the screen). Image content was defined by 30 semantic component scores derived from the 300-dimensional Word2Vec vectors from the COCO annotations (**Fig. 2**). The 30 semantic components explained 68.59% of the variance in COCO annotations for the images used in the study. As with the behavioral analyses above, we assessed reconstruction accuracy by comparing within-item vs. across-item similarity. Here, however, within-item similarity was defined as the Fisher-transformed Pearson correlation between the reconstructed component scores for a given image (as predicted from the inverted fMRI encoding model) and the ‘actual’ semantic component scores for that image (derived from COCO annotations). Across-item similarity was defined as the mean Fisher-transformed Pearson correlation between predicted component scores for a given image and actual component scores for *different images* (from the same session). Reconstruction of content information was determined to be successful if within-item similarity was greater than across-item similarity, as determined by mixed-effects linear models which included subject and session numbers as random factors. Consistent with our previous studies (Cowen et al. 2014; Lee and Kuhl 2016), robust reconstruction accuracies were obtained from visual regions (VTC: χ ^2^_1_ = 3024.1, *p* < 0.0001, β = 0.136, SE = 0.002; OTC: χ ^2^_1_ = 3785.6, *p* < 0.0001, β = 0.146, SE = 0.002) as well as ANG and other lateral parietal ROIs (ANG: χ ^2^_1_ = 754.4, β = 0.063, SE = 0.002; SMG: χ ^2^_1_ = 277.8, β = 0.040, SE = 0.002; SPL: χ ^2^_1_ = 624.0, β = 0.059, SE = 0.002; IPS: χ ^2^_1_ = 774.5, β = 0.066, SE = 0.002; *p* values < 0.0001) (**Fig. 4A**,**B**). However, reconstruction accuracies sharply varied across ROIs (main effect of ROI from repeated-measures ANOVA: *F*_5,90_ = 350.69, *p* < 0.0001, *η* ^2^_p_= 0.95), with higher accuracies in the visual ROIs compared to the parietal ROIs (*p*’s < 0.0001 for all paired-samples *t*-tests comparing the VTC and OTC ROIs to each of the lateral parietal ROIs).

### 3.3. Reconstruction of image content from cued recall task

Next, we extended our method to test for content reconstruction for images retrieved from memory during the cued recall task. Critically, and in contrast to recognition-based reconstructions for which the to-be-reconstructed image was visually present, here the to-be-reconstructed image was visually absent (subjects were only shown the texture cues) thus requiring top-down retrieval of the target image. For this analysis, we trained the semantic encoding model with ‘old’ images from the recognition set (exploiting the large number of recognition trials) but tested it on images from the cued recall task. As with the recognition-based reconstructions, evidence for successful recall-based reconstructions was obtained if within-item similarity (correlations between the semantic component scores predicted from the inverted encoding model and the ‘target’ semantic component scores) were reliably higher than across-item correlations. As described in the following sections, we used several approaches for defining the ‘target’ semantic component scores.

As a first step, we defined target semantic component scores based on the COCO annotations (as in the preceding section which tested for reconstruction accuracy during the recognition memory task). Successful recall-based content reconstruction was observed across each of the visual and parietal regions (VTC: χ ^2^_1_ = 83.9, *p* < 0.0001, β = 0.083, SE = 0.009; OTC: χ ^2^_1_ = 24.6, *p* < 0.0001, β = 0.043, SE = 0.009; ANG: χ ^2^_1_ = 26.6, *p* < 0.0001, β = 0.049, SE = 0.009; SMG: χ ^2^_1_ = 26.0, *p* = 0.0001, β = 0.048, SE = 0.009; SPL: χ ^2^_1_ = 24.5, *p* < 0.0001, β = 0.047, SE = 0.009; IPS: χ ^2^_1_ = 32.6, *p* < 0.0001, β = 0.056, SE = 0.010; **Fig. 4C**). Accuracy significantly varied across ROIs (main effect of ROI: *F*_5,90_ = 3.46, *p* = 0.007, *η* ^2^_p_ = 0.16), with accuracy numerically highest in VTC. To provide a sense of the subjective accuracy of recall-based reconstructions, **Fig. 5** shows examples of words that were most similar to the reconstructed semantic components (generated using the “most_similar” function of Word2Vec) for images with varying degrees of recall-based reconstruction accuracy. Specifically, we pooled the reconstructed semantic component scores in VTC across all subjects and sessions, and then rank ordered these reconstructed scores by accuracy (match to the target scores). **Fig. 5** shows examples of the “most similar” words for reconstructions that were in the top 1%, top 25%, and top 50%.

**Fig. 5.**
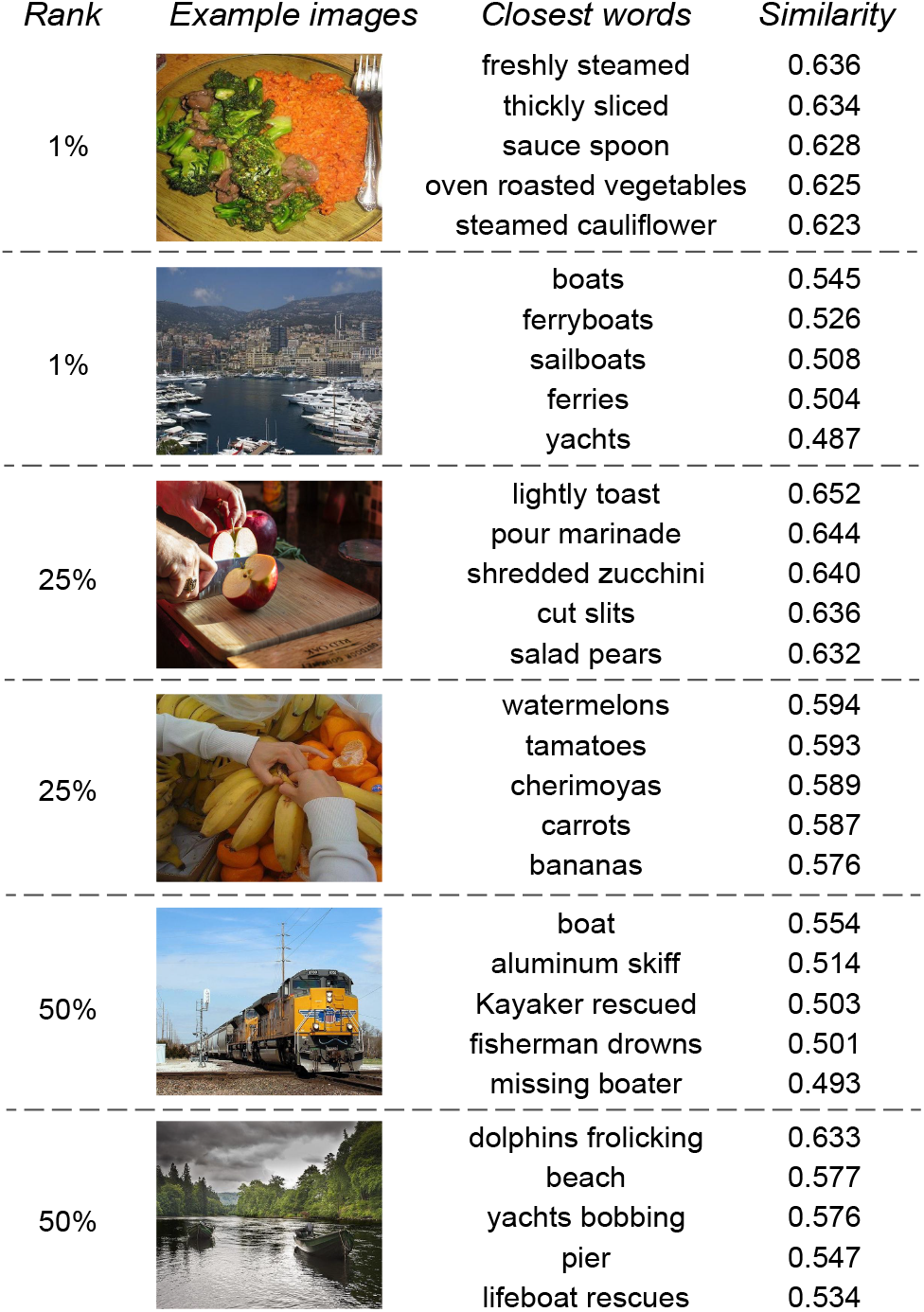
Examples of reconstructed image content from VTC. (Left) Rank of the reconstruction accuracy pooled over all subjects and sessions. (Middle left) Example images being recalled. (Middle right) The top 5 most similar words and word combinations describing the semantic component scores reconstructed from VTC. The words were generated by the Word2Vec default ‘most_similar’ function. (Right) Similarity scores between vectors corresponding to the content reconstructed from VTC and vectors of the Word2Vec most similar words.

While all ROIs exhibited above-chance content reconstruction for both recognition-based and recall-based reconstructions, the difference between recognition-versus recall-based reconstructions markedly varied across ROIs, as reflected by an interaction between trial type (recognition, recall) and ROI (*F*_5,90_ = 24.88, *p* < 0.0001, *η* ^2^_p_= 0.58). Whereas content reconstruction accuracy was much higher for recognition than recall in VTC (χ ^2^_1_ = 15.6, *p* < 0.0001, β = 0.052, SE = 0.012) and OTC (χ ^2^_1_ = 50.7, *p* < 0.0001, β = 0.103, SE = 0.010), reconstruction accuracy in parietal regions did not significantly differ for recognition versus recall trials (ANG: χ ^2^_1_ = 2.16, β = 0.014, SE = 0.009; SMG: χ ^2^_1_ = 0.58, β = -0.008, SE = 0.011; SPL: χ ^2^_1_ = 1.49, β = 0.012, SE = 0.010; IPS: χ ^2^_1_ = 0.74, β = 0.010, SE = 0.011; *p* values > 0.140) (**Fig. 4D**). Thus, whereas VTC and OTC exhibited a clear ‘preference’ for images that were visually present (recognition trials), reconstructions from parietal regions were of comparable success when images were visually present (recognition trials) or entirely driven by memory (recall trials).

### 3.4. Similarity between reconstructed content and verbal descriptions of memories

In the preceding analyses, the target semantic content of each image was defined by image annotations that are part of the COCO image dataset. We next tested the degree to which semantic component scores reconstructed from the inverted fMRI encoding models (measured during the scanned cued recall task) matched the semantic component scores derived from subjects’ *own verbal memory* of each image (measured during the post test) (**Fig. 6A**). As described for behavioral analysis of the verbal recall data (**Fig. 3C**), each subject’s verbal recall of each image was translated into 30 semantic component scores. These target component scores could then be readily compared to (correlated with) the semantic component scores predicted from the inverted fMRI encoding models. Again, we found higher within-than across-item correlations in each of the visual and parietal ROIs (**Fig. 6B**) (VTC: χ ^2^_1_ = 49.7, *p* < 0.0001, β = 0.087, SE = 0.012; OTC: χ ^2^_1_ = 21.2, *p* < 0.0001, β = 0.055, SE = 0.011; ANG: χ ^2^_1_ = 14.1, *p* < 0.0001, β = 0.050, SE = 0.013; SMG: χ ^2^_1_ = 12.3, *p* = 0.0004, β = 0.047, SE = 0.001; SPL: χ ^2^_1_ = 7.4, *p* = 0.007, β = 0.038, SE = 0.014; IPS: χ ^2^_1_ = 7.4, *p* = 0.007, β = 0.036, SE = 0.013). Accuracy varied across ROIs (main effect of ROI: *F*_5,90_ = 2.75, *p* = 0.024, *η* ^2^_p_= 0.13), with accuracy numerically highest in VTC. These results confirm that the reconstructed semantic information from LPC and visual regions matched subjects’ verbal descriptions of their memories.

**Fig. 6.**
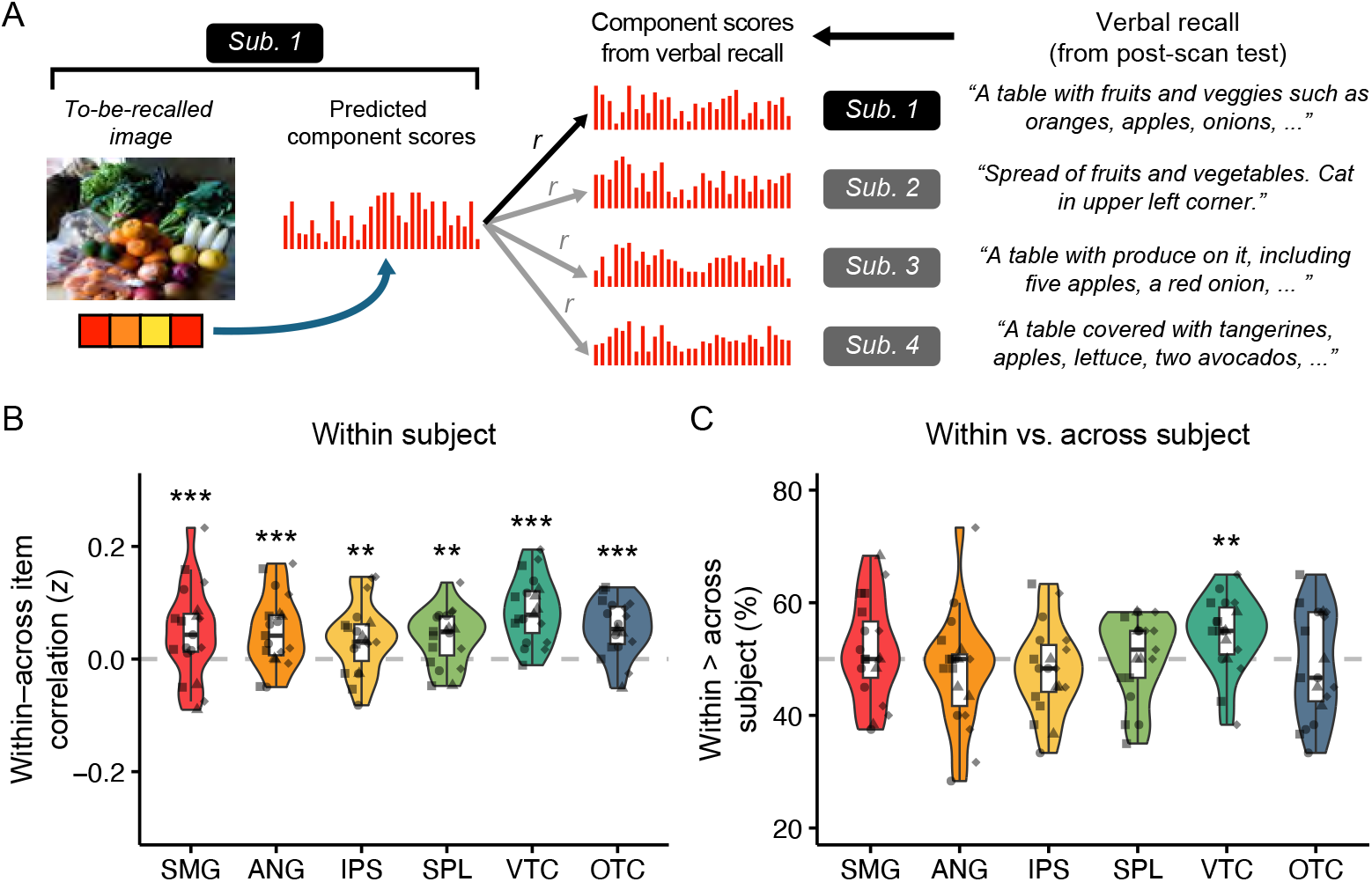
Correspondence between semantic component scores reconstructed from fMRI vs. derived from verbal recall. **A**. Schematic of the analysis. For each to-be-recalled image for each subject, semantic component scores were reconstructed (predicted) from fMRI activity patterns using semantic encoding models trained on the recognition trials and tested on the recall trials. These reconstructed semantic component scores were then correlated with semantic component scores derived from subjects’ verbal recall of the same image (measured during the post-scan overt cued recall test). **B**. Reconstruction accuracy as reflected by the difference between within-item vs. across-item correlations, with all correlations performed within-subject. Reconstruction accuracy was significantly above chance for all ROIs. **C**. Subject-specific reconstructions. To test for subject-specific (idiosyncratic) reconstructions, the semantic component scores reconstructed from one subject’s fMRI data were correlated with semantic component scores generated from (i) the same subject’s verbal recall data (e.g., Sub. 1 -> Sub. 1, black arrow, in **A**) and (ii) other subjects’ verbal recall data of the exact same images (e.g., Sub-1 -> Sub. 2, grey arrows, in **A**). Reconstructions were considered to contain subject-specific information within-subject correlations were higher than the across-subject correlations. Data shown reflect the mean percentage of within-subject correlations that exceeded across-subject correlations. Accuracy was significantly above chance (dash line, 50%) only for VTC. Notes: dots represent data from individual sessions with each subject represented by a different shape; ** *p* < 0.01, *** *p* < 0.001.

While the preceding analysis confirms a match between verbal recall and reconstructed semantic component scores, an even stricter test is whether the semantic component scores reconstructed from a given subject’s fMRI data more closely resembled the semantic component scores from *that subject’s verbal recall* compared to semantic component scores from other subjects’ verbal recall of the exact same images. To test this, we first calculated the Pearson correlations between the semantic component scores reconstructed from a given subject’s inverted fMRI encoding model and the corresponding semantic component scores derived from that same subject’s verbal recall (within-subject similarity). We then compared this within-subject similarity to across-subject similarity: the correlations between a given subject’s reconstructed semantic component scores and the corresponding semantic component scores derived from *different subjects’* verbal recall of the same images. It is important to emphasize that both of these measures were *within-item* correlations (i.e., they relate to the exact same images). If within-subject similarity exceeds across-subject similarity, this provides evidence for a subject-specific correspondence between fMRI-based reconstructions and verbal recall.

For each subject, session, and ROI we compared within-subject similarity to across-subject similarity in order to generate an accuracy score for image. This image-specific accuracy score reflected the percentage of comparisons for which within-subject correlations were greater than across-subject correlations. For example, for a given image recalled by subject 1, the fMRI-based reconstructed semantic component scores would be correlated with the semantic component scores derived from verbal recall from subject 1 (within-subject similarity) and with the semantic component scores derived from verbal recall from subjects 2, 3 and 4 (across-subject similarity). If, for example, the within-subject correlation [*r*(1,1)] was greater than two of the three possible across-subject correlations [*r*(1,2), *r*(1,3), *r*(1,4)], this would correspond to an accuracy of 66.66% for that image. In this manner, the mean accuracy was computed for each subject, session, and ROI. Chance-level accuracy was 50% (i.e., by chance, within-subject similarity should exceed across-subject similarity 50% of the time). Strikingly, we observed above-chance accuracy—i.e., subject-specific reconstructions—in VTC (54.39%, *t*(18) = 2.90, *p* = 0.009, Cohen’s d = 0.66; **Fig. 6B**)—which was also the ROI that exhibited the highest recall-based reconstruction accuracy in each of the preceding analyses. Accuracy did not exceed chance in any of the other ROIs [OTC: *M* = 49.21%, *t*(18) = -0.36, *p* = 0.720, Cohen’s d = -0.08; ANG: *M =* 47.76%, *t*(18) = -0.96, *p* = 0.348, Cohen’s d = -0.22; SMG: *M* = 51.36%, *t*(18) = 0.67, *p* = 0.512, Cohen’s d = 0.20; SPL: *M* = 50.40%, t(18) = 0.24, *p* = 0.816, Cohen’s d = 0.05; IPS: *M* = 48.11%, *t*(18) = -1.04, *p* = 0.310, Cohen’s d = -0.24; **Fig. 6B**].

### 3.5. Across-subject reconstruction of recalled memories

Finally, we tested whether information ‘learned’ by the semantic encoding models (i.e., the mappings between voxel activity patterns and semantic component scores) successfully transferred across individuals. More specifically, we tested whether the contents of memory recall for each subject could be reconstructed using encoding models trained on data from independent subjects. To test this, we iteratively trained semantic encoding models using the recognition data from three of the four subjects and tested the model on recall trials from the held-out subject. That is, the weight matrix that was applied to each subject’s fMRI activity patterns from the recall trials was entirely derived from independent subjects. We first tested content reconstruction accuracy by correlating the reconstructed component scores with component scores derived from the COCO annotations (as in **Fig. 4C**). Again, within-item similarity was compared against across-item similarity. Successful reconstruction (greater within-item similarity than across-item similarity) was observed in ANG (χ ^2^_1_ = 13.6, *p* = 0.0002, β = 0.033, SE = 0.009), SPL (χ ^2^_1_ = 5.5, *p* = 0.020, β = 0.022, SE = 0.010), IPS (χ ^2^_1_ = 5.4, *p* = 0.020, β = 0.022, SE = 0.009), VTC (χ ^2^_1_ = 37.8, *p* < 0.0001, β = 0.056, SE = 0.009), and OTC (χ ^2^_1_ = 10.5, *p* = 0.001, β = 0.027, SE = 0.008) (**Fig. 7A**).

**Fig. 7.**
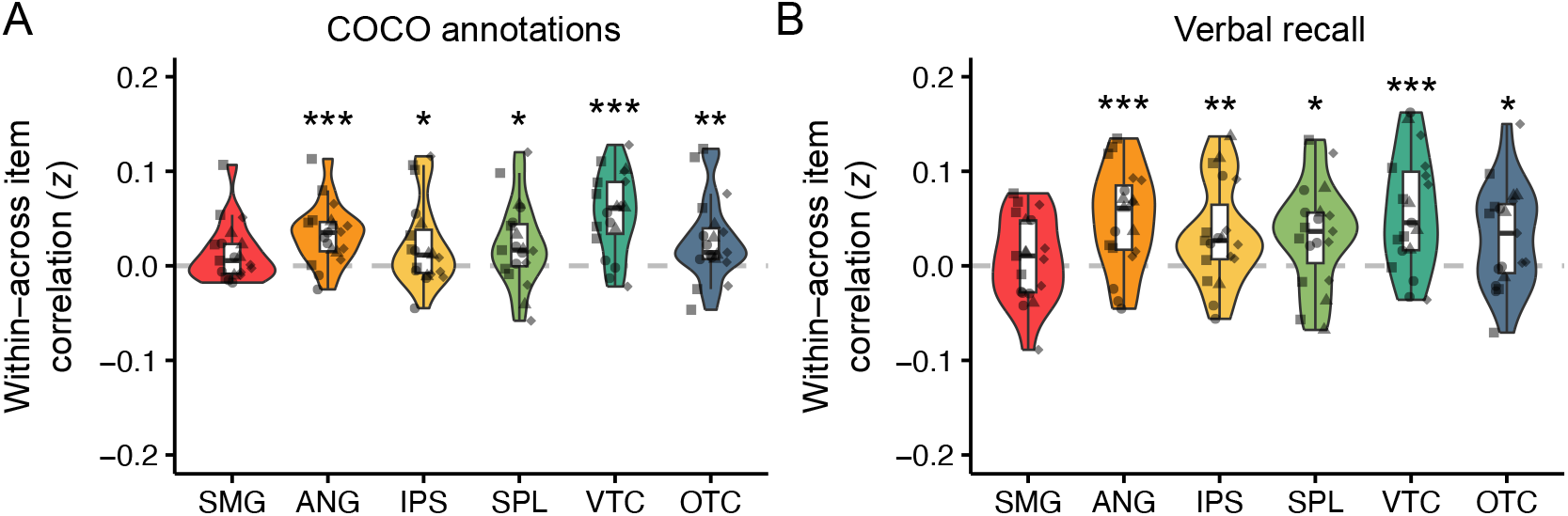
Across-subject application of the semantic encoding models. For these analyses, the semantic encoding model was iteratively trained on recognition trials from 3 of the 4 subjects and then tested on recall trials from the held-out subject. **A**. Mean accuracy of reconstructed semantic component scores for each ROI based on comparison to semantic component scores derived from COCO annotations (within-item correlations – across-item correlations). **B**. Mean reconstruction accuracy for each ROI based on comparison to semantic component scores derived from verbal recall (within-item correlations – across-item correlations). For **B**, although the training/testing of the encoding models was performed across subjects, the covert recall trials used for reconstructing the semantic component scores and the verbal recall trials used for testing accuracy were always within the same subject. Notes: dots represent data from individual sessions with each subject represented by a different shape; ** *p* < 0.05, ** *p* < 0.01, *** *p* < 0.001, two tailed.

We next replicated this analysis with the only difference being that reconstructed component scores were correlated with component scores derived from each subject’s (own) verbal recall (as in **Fig. 6B**). Again, within-item similarity was greater than across-item similarity in ANG (χ ^2^_1_ = 15.7, *p* < 0.0001, β = 0.049, SE = 0.012), SPL (χ ^2^_1_ = 6.4, *p* = 0.011, β = 0.032, SE = 0.013), IPS (χ ^2^_1_ = 7.7, *p* = 0.005, β = 0.034, SE = 0.012), VTC (χ ^2^_1_ = 24.1, *p* < 0.0001, β = 0.056, SE = 0.011), and OTC (χ ^2^_1_ = 6.4, *p* = 0.011, β = 0.031, SE = 0.012) (**Fig. 7B**). These findings provide evidence that, across subjects, the mappings between semantic content and fMRI activity patterns were shared to a degree that encoding models could be transferred to independent subjects to reconstruct the contents of memory recall.

## 4. Discussion

In the current study, we extracted high-level semantic features from complex natural images and modeled relationships between these semantic features and fMRI activity patterns using voxelwise encoding models. By inverting the encoding models, we tested whether the semantic content of retrieved memories could be reconstructed from evoked fMRI activity patterns. Using a multiple-session training procedure, we show that semantic content was successfully reconstructed from fMRI activity patterns in lateral parietal and visual cortices. Notably, however, reconstruction accuracy differed across these regions according to whether images were visually present (during recognition) or cued by arbitrarily-associated abstract images (during recall). Whereas reconstruction accuracy in visual cortex was markedly lower when images were recalled from memory (recall trials) compared to when they were visually present (recognition trials), lateral parietal regions were relatively insensitive to this difference between trial types. Separately, by applying natural language processing methods to subjects’ verbal recall data and projecting these recall data into the same feature space as the fMRI reconstructions, we also established that fMRI-based reconstructions reliably matched subjects’ verbal recall data. In fact, reconstructions from ventral temporal cortex reflected idiosyncratic differences in how different subjects remembered the exact same image. Finally, we show that encoding models trained on a subset of subjects reliably transferred to held-out subjects, indicating that the mapping between fMRI activity patterns and semantic content was consistent enough across subjects to allow for across-subject reconstructions. Collectively, these findings provide important evidence for multidimensional memory representations in lateral parietal and visual cortices and establish the relevance of these neural representations to complex behavioral expressions of memory recall.

### 4.1. Reconstruction and recall of multidimensional memory representations

Numerous prior fMRI studies have demonstrated content-sensitivity of fMRI activity patterns in visual and lateral parietal cortices during memory retrieval (Favila et al. 2018; Kuhl et al. 2011; Kuhl and Chun 2014; Lee et al. 2019; Polyn 2005; St-Laurent et al. 2015). However, the majority of this evidence comes from studies that have measured an objective, single stimulus property or dimension. For example, many studies have tested for decoding of visual category information (Kuhl et al. 2011; Polyn 2005). Others have demonstrated an item-specific ‘match’ between fMRI activity patterns elicited during memory encoding and those elicited during memory retrieval (Favila et al. 2018; Kuhl and Chun 2014; Lee et al. 2019; St-Laurent et al. 2015). While the current findings also constitute evidence for item-specific representations (in that our analyses revealed differences between individual scene images), the key difference in the current study is that item-specific representations were ‘built’ by predicting and combining constituent features (Lee and Kuhl 2016; Naselaris et al. 2011). In fact, reconstructions were based on encoding models that were not trained on the to-be-reconstructed images (Brouwer and Heeger 2009). Thus, the stimulus-specific representations observed here cannot be explained by subjects generating verbal labels or stimulus-specific tags during encoding and then re-expressing that label/tag during recall.

The motivation for establishing multidimensional neural representations of memories is that these measures have the potential to capture the richness, subjectivity, and idioscynracies with which real world memories are recalled. Critically, however, validation of these neural representations requires behavioral expressions of memory that also capture the same richness, subjectivity, and idiosyncracies. Our solution to this problem was to use natural language processing methods that allowed our fMRI and behavioral data to be described using the same feature dimensions. Considering the behavioral recall data alone, text embeddings were highly sensitive to differences between images (**Fig. 3C**) validating the use of this method to characterize verbal recall data (Heusser et al. 2021; Song, Finn, and Rosenberg 2021). Moreover, across visual and lateral parietal ROIs, there was strong correspondence between fMRI-based reconstructions and subjects’ verbal recall (**Fig. 6B**), demonstrating that the multidimensional fMRI reconstructions aligned with the multidimensional expressions of verbal recall. Most strikingly, reconstructions generated from ventral temporal cortex were significantly more similar to subjects’ own verbal recall compared to other subjects’ verbal recall of exactly the same images (**Fig. 6C**). In other words, ventral temporal cortex reconstructions reflected subjective or idiosyncratic differences in how scene images were remembered. This effect is particularly notable when considering that there were no experimental pressures for subjects to use unique language or to differentiate their responses from other subjects. Thus, these methods may be even more sensitive to subjective/idiosyncratic information in experimental contexts where there are factors that promote memory differentiation (Favila, Chanales, and Kuhl 2016; Hulbert and Norman 2015; Kim, Norman, and Turk-Browne 2017).

### 4.2. Reconstructions in lateral parietal cortex versus visual cortical areas

Not surprisingly, reconstructions from visual cortical areas (VTC, OTC) were markedly higher when images were visually present (recognition trials) compared to when they were visually absent (recall trials). In contrast, this fundamental distinction between trial types did not significantly influence reconstruction accuracy in LPC regions. Notably, several recent studies have specifically shown that, in contrast to visual cortical regions, LPC representations are stronger during memory recall compared to memory encoding or perception (Akrami et al. 2018; Favila et al. 2018, 2020; Long and Kuhl 2021; Xiao et al. 2017). While a definitive account of why LPC is biased towards memory-based information is not yet clear (Favila et al. 2020), the current findings provide additional support for a relative preference toward memory-based information in LPC. Here, however, we did not observe *stronger* (more accurate) LPC reconstructions during recall compared to recognition. That said, it is important to emphasize that recognition-based reconstructions were generated from models trained and tested on recognition trials whereas recall-based reconstructions were generated from models trained on recognition trials but *tested on recall trials*. Thus, a direct comparison of reconstruction accuracy for recall versus recognition trials is not an apples-to-apples comparison. Instead, the critical statistical comparison is the *relative* sensitivity of visual versus LPC regions to the difference in trial types. Indeed, this interaction was highly significant (**Fig. 4D**).

An obvious question raised by the current findings is whether recall reconstructions would be significantly better (in LPC and possibly VTC, as well) if the encoding model had been trained only on recall trials (Chen et al. 2017). In our study, this was not feasible because the number of recall trials was relatively small (far fewer than the number of recognition trials). At a practical level, recall trials are also much harder to include in large numbers because they depend on pre-training the paired associations (e.g., we used an extensive training procedure to ensure successful, vivid recall; **Fig. 1**). However, in an effort to address the potential concern of poor transfer from ‘pure perception’ trials to recall trials, we opted to pre-expose subjects to images in the recognition set such that the images used for model training were ‘old’ images. The sole rationale for the pre-exposure phase was that the semantic encoding models might better transfer to recall trials if the training trials had some memory component. Specifically, we reasoned that the representational format of a recall trial might be more similar to an ‘old’ recognition trial than to an entirely novel stimulus. While this thinking was informed by recent findings (Akrami et al. 2018; Favila et al. 2018, 2020; Long and Kuhl 2021; Xiao et al. 2017), it was not our intention—nor are we able—to test whether this design feature actually improved model transfer. That said, it does represent an interesting question that could be tested empirically in future studies.

While we observed evidence for idiosyncratic (subject-specific) relationships between fMRI-based reconstructions and verbal recall when considering reconstructions from VTC, we did not observe significant relationships for any of the LPC ROIs. On the one hand, this null result for LPC regions is surprising in light of evidence that memory reactivation in LPC has been associated with subjective qualities of memory recall (Bone et al. 2020; Johnson et al. 2015; Kuhl and Chun 2014; Richter et al. 2016). However, across analyses, reconstruction accuracy was higher in VTC than in LPC ROIs, meaning there simply may have been better sensitivity within VTC to subtle differences in within-subject versus across-subject comparisons. As described above, it is possible that training the encoding models on recall trials (as opposed to recognition trials) might boost performance in LPC ROIs and thereby improve sensitivity to subject-specific differences. Indeed, we view this as a very interesting and reasonable possibility. Alternatively, it is possible that LPC preferentially expresses representational formats of retrieved memories that are relatively shared across subjects (Chen et al., 2018). Given that both of these are viable possibilities, we would caution against drawing conclusions based on the absence of significant subject-specific effects in the LPC ROIs. Instead, we view the significant results in VTC as a proof of concept that our methodological approach can be used to identify subject-specific idiosyncrasies in how complex images are remembered.

### 4.3. Semantic encoding models generalize across subjects

Although we deliberately used an extensive-sampling procedure to maximize the amount of within-subject training data available for the encoding models, we also show that encoding models transferred quite well across subjects. Specifically, training encoding models using recognition trials from N-1 subjects allowed for successful recall-based reconstruction in held out subjects (**Fig. 7**). This successful transfer across subjects indicates that the mapping between semantic components and fMRI activity patterns was shared— at least to some degree—across different individuals. Importantly, this shared mapping between semantic information and fMRI activity patterns is not at odds with our finding (or the idea) of idiosyncratic memory representations. For example, consider two individuals that had breakfast together. These individuals may have a common neural representation of the concept of coffee, and each of them may have had coffee for breakfast. However, when remembering breakfast, these individuals may differ in the degree to which the concept of coffee is a salient component of their memory and, therefore, in the degree to which the neural representation of coffee is activated when they remember breakfast. Thus, leveraging shared mappings (i.e., encoding models trained across different individuals) need not come at the expense of identifying idiosyncratic ways in which individuals perceive or remember their experiences (Finn et al. 2018, 2020).

More generally, the success of the across-subject encoding models has two main implications. First, this finding adds to a growing body of evidence that, even for complex and naturalistic stimuli (e.g., movies and narratives), there is a surprising degree of consistency across individuals in how these stimuli are represented in patterns of neural activity (Chen et al. 2017; Finn et al. 2018; Hasson et al. 2004; Zadbood et al. 2017). Second, leveraging across-subject encoding models could have substantial practical—and theoretical—advantages. For example, as noted above, it was not feasible in our experimental paradigm for each subject to learn and recall thousands of different scenes (due to the training time it would require and the deterioration in memory performance that would be expected with such a large memory set). However, it is much more feasible to obtain thousands of recall trials *across subjects*. Thus, some analyses which are impractical—or that would be data starved—within subjects, might become feasible if across-subject models are leveraged. Moreover, a single well-powered training data set could potentially be applied to many distinct test sets. Finally, it is also notable that here, we only aligned across-subject data in anatomical space. Additional gains in across-subject transfer may well be realized by aligning data in a common high-dimensional functional space (Chen et al. 2015; Haxby et al. 2011, 2020).

### 4.4. Conclusions

To summarize, we used inverted semantic encoding models applied to fMRI data to reconstruct multidimensional content in natural scene images as they were viewed and recalled from memory. We found that visual and lateral parietal cortices supported successful reconstructions both when viewing and recalling images. However, whereas lateral parietal reconstructions were relatively insensitive to whether images were viewed or recalled from memory, visual cortical reconstructions were markedly lower for recalled versus viewed images. Additionally, by applying natural language processing methods to behavioral measures of memory recall, we show that fMRI-based reconstructions of recalled content matched subjects’ verbal recall and that fMRI-based reconstructions even reflected idiosyncratic qualities of subjects’ recall. Finally, we show that semantic encoding models reliably transferred across individuals, allowing for successful reconstruction of a given subject’s memory using encoding models trained on entirely different individuals. Collectively, these findings provide important evidence characterizing multidimensional memory representations and linking their neural and behavioral expressions.

## Acknowledgements

The research described here was supported by NSF CAREER Award BCS-1752921 and NIH-NINDS R01 NS107727 to B.A.K.

## References

Akrami, Athena, Charles D. Kopec, Mathew E. Diamond, and Carlos D. Brody. 2018. “Posterior Parietal Cortex Represents Sensory History and Mediates Its Effects on Behaviour.” Nature 554(7692):368–72. doi: 10.1038/nature25510.

Bone, Michael B., Fahad Ahmad, and Bradley R. Buchsbaum. 2020. “Feature-Specific Neural Reactivation during Episodic Memory.” Nature Communications 11(1):1945. doi: 10.1038/s41467-020-15763-2.

Bonnici, H. M., F. R. Richter, Y. Yazar, and J. S. Simons. 2016. “Multimodal Feature Integration in the Angular Gyrus during Episodic and Semantic Retrieval.” Journal of Neuroscience 36(20):5462–71. doi: 10.1523/JNEUROSCI.4310-15.2016.

Brouwer, G. J., and D. J. Heeger. 2009. “Decoding and Reconstructing Color from Responses in Human Visual Cortex.” Journal of Neuroscience 29(44):13992–3. doi: 10.1523/JNEUROSCI.3577-09.2009.

Chen, Janice, Yuan Chang Leong, Christopher J. Honey, Chung H. Yong, Kenneth A. Norman, and Uri Hasson. 2017. “Shared Memories Reveal Shared Structure in Neural Activity across Individuals.” Nature Neuroscience 20(1):115–25. doi: 10.1038/nn.4450.

Chen, Po-Hsuan (Cameron), Janice Chen, Yaara Yeshurun, Uri Hasson, James Haxby, and Peter J. Ramadge. 2015. “A Reduced-Dimension FMRI Shared Response Model.” in Advances in Neural Information Processing Systems. Vol. 28, edited by C. Cortes, N. Lawrence, D. Lee, M. Sugiyama, and R. Garnett. Curran Associates, Inc.

Cooper, Rose A., and Maureen Ritchey. 2019. “Cortico-Hippocampal Network Connections Support the Multidimensional Quality of Episodic Memory.” ELife 8:e45591. doi: 10.7554/eLife.45591.

Cowen, Alan S., Marvin M. Chun, and Brice A. Kuhl. 2014. “Neural Portraits of Perception: Reconstructing Face Images from Evoked Brain Activity.” NeuroImage 94:12–22. doi: 10.1016/j.neuroimage.2014.03.018.

Danker, Jared F., and John R. Anderson. 2010. “The Ghosts of Brain States Past: Remembering Reactivates the Brain Regions Engaged during Encoding.” Psychological Bulletin 136(1):87–102. doi: 10.1037/a0017937.

Esteban, Oscar, Daniel Birman, Marie Schaer, Oluwasanmi O. Koyejo, Russell A. Poldrack, and Krzysztof J. Gorgolewski. 2017. “MRIQC: Advancing the Automatic Prediction of Image Quality in MRI from Unseen Sites.” PLOS ONE 12(9):e0184661. doi: 10.1371/journal.pone.0184661.

Esteban, Oscar, Christopher J. Markiewicz, Ross W. Blair, Craig A. Moodie, A. Ilkay Isik, Asier Erramuzpe, James D. Kent, Mathias Goncalves, Elizabeth DuPre, Madeleine Snyder, Hiroyuki Oya, Satrajit S. Ghosh, Jessey Wright, Joke Durnez, Russell A. Poldrack, and Krzysztof J. Gorgolewski. 2019. “FMRIPrep: A Robust Preprocessing Pipeline for Functional MRI.” Nature Methods 16(1):111–16. doi: 10.1038/s41592-018-0235-4.

Ester, Edward F., Thomas C. Sprague, and John T. Serences. 2015. “Parietal and Frontal Cortex Encode Stimulus-Specific Mnemonic Representations during Visual Working Memory.” Neuron 87(4):893– 905. doi: 10.1016/j.neuron.2015.07.013.

Favila, Serra E., Avi J. H. Chanales, and Brice A. Kuhl. 2016. “Experience-Dependent Hippocampal Pattern Differentiation Prevents Interference during Subsequent Learning.” Nature Communications 7(1). doi: 10.1038/ncomms11066.

Favila, Serra E., Hongmi Lee, and Brice A. Kuhl. 2020. “Transforming the Concept of Memory Reactivation.” Trends in Neurosciences S0166223620302137. doi: 10.1016/j.tins.2020.09.006.

Finn, Emily S., Philip R. Corlett, Gang Chen, Peter A. Bandettini, and R. Todd Constable. 2018. “Trait Paranoia Shapes Inter-Subject Synchrony in Brain Activity during an Ambiguous Social Narrative.” Nature Communications 9(1):2043. doi: 10.1038/s41467-018-04387-2.

Finn, Emily S., Enrico Glerean, Arman Y. Khojandi, Dylan Nielson, Peter J. Molfese, Daniel A. Handwerker, and Peter A. Bandettini. 2020. “Idiosynchrony: From Shared Responses to Individual Differences during Naturalistic Neuroimaging.” NeuroImage 215:116828. doi: 10.1016/j.neuroimage.2020.116828.

Gilmore, Adrian W., Steven M. Nelson, and Kathleen B. McDermott. 2015. “A Parietal Memory Network Revealed by Multiple MRI Methods.” Trends in Cognitive Sciences 19(9):534–43. doi: 10.1016/j.tics.2015.07.004.

Gilmore, Adrian W., Alina Quach, Sarah E. Kalinowski, Stephen J. Gotts, Daniel L. Schacter, and Alex Martin. 2021. “Dynamic Content Reactivation Supports Naturalistic Autobiographical Recall in Humans.” The Journal of Neuroscience 41(1):153–66. doi: 10.1523/JNEUROSCI.1490-20.2020.

Hasson, Uri, Yuval Nir, Ifat Levy, Galit Fuhrmann, and Rafael Malach. 2004. “Intersubject Synchronization of Cortical Activity During Natural Vision.” Science 303(5664):1634–40. doi: 10.1126/science.1089506.

Haxby, James V., J. Swaroop Guntupalli, Andrew C. Connolly, Yaroslav O. Halchenko, Bryan R. Conroy, M. Ida Gobbini, Michael Hanke, and Peter J. Ramadge. 2011. “A Common, High-Dimensional Model of the Representational Space in Human Ventral Temporal Cortex.” Neuron 72(2):404–16. doi: 10.1016/j.neuron.2011.08.026.

Haxby, James V., J. Swaroop Guntupalli, Samuel A. Nastase, and Ma Feilong. 2020. “Hyperalignment: Modeling Shared Information Encoded in Idiosyncratic Cortical Topographies.” ELife 9:e56601. doi: 10.7554/eLife.56601.

Heusser, Andrew C., Paxton C. Fitzpatrick, and Jeremy R. Manning. 2021. “Geometric Models Reveal Behavioural and Neural Signatures of Transforming Experiences into Memories.” Nature Human Behaviour. doi: 10.1038/s41562-021-01051-6.

Hulbert, J. C., and K. A. Norman. 2015. “Neural Differentiation Tracks Improved Recall of Competing Memories Following Interleaved Study and Retrieval Practice.” Cerebral Cortex 25(10):3994–4008. doi: 10.1093/cercor/bhu284.

Huth, Alexander G., Wendy A. de Heer, Thomas L. Griffiths, Frédéric E. Theunissen, and Jack L. Gallant. 2016. “Natural Speech Reveals the Semantic Maps That Tile Human Cerebral Cortex.” Nature 532(7600):453–58. doi: 10.1038/nature17637.

Johnson, Marcia K., Brice A. Kuhl, Karen J. Mitchell, Elizabeth Ankudowich, and Kelly A. Durbin. 2015. “Age-Related Differences in the Neural Basis of the Subjective Vividness of Memories: Evidence from Multivoxel Pattern Classification.” Cognitive, Affective, & Behavioral Neuroscience 15(3):644– 61. doi: 10.3758/s13415-015-0352-9.

Kay, Kendrick N., Thomas Naselaris, Ryan J. Prenger, and Jack L. Gallant. 2008. “Identifying Natural Images from Human Brain Activity.” Nature 452(7185):352–55. doi: 10.1038/nature06713.

Kim, Ghootae, Kenneth A. Norman, and Nicholas B. Turk-Browne. 2017. “Neural Differentiation of Incorrectly Predicted Memories.” The Journal of Neuroscience 37(8):2022–31. doi: 10.1523/JNEUROSCI.3272-16.2017.

Kok, Peter, Lindsay I. Rait, and Nicholas B. Turk-Browne. 2020. “Content-Based Dissociation of Hippocampal Involvement in Prediction.” Journal of Cognitive Neuroscience 32(3):527–45. doi: 10.1162/jocn_a_01509.

Kuhl, B. A., and M. M. Chun. 2014. “Successful Remembering Elicits Event-Specific Activity Patterns in Lateral Parietal Cortex.” Journal of Neuroscience 34(23):8051–60. doi: 10.1523/JNEUROSCI.4328-13.2014.

Kuhl, B. A., J. Rissman, M. M. Chun, and A. D. Wagner. 2011. “Fidelity of Neural Reactivation Reveals Competition between Memories.” Proceedings of the National Academy of Sciences 108(14):5903–8. doi: 10.1073/pnas.1016939108.

Lee, Hongmi, and Brice A. Kuhl. 2016. “Reconstructing Perceived and Retrieved Faces from Activity Patterns in Lateral Parietal Cortex.” The Journal of Neuroscience 36(22):6069–82. doi: 10.1523/JNEUROSCI.4286-15.2016.

Lee, Hongmi, Rosalie Samide, Franziska R. Richter, and Brice A. Kuhl. 2019. “Decomposing Parietal Memory Reactivation to Predict Consequences of Remembering.” Cerebral Cortex 29(8):3305–18. doi: 10.1093/cercor/bhy200.

Lin, Tsung-Yi, Michael Maire, Serge Belongie, Lubomir Bourdev, Ross Girshick, James Hays, Pietro Perona, Deva Ramanan, C. Lawrence Zitnick, and Piotr Dollár. 2015. “Microsoft COCO: Common Objects in Context.” ArXiv:1405.0312 [Cs].

Long, Nicole M., and Brice A. Kuhl. 2021. “Cortical Representations of Visual Stimuli Shift Locations with Changes in Memory States.” Current Biology 31(5):1119-1126.e5. doi: 10.1016/j.cub.2021.01.004.

Naselaris, Thomas, Emily Allen, and Kendrick Kay. 2021. “Extensive Sampling for Complete Models of Individual Brains.” Current Opinion in Behavioral Sciences 40:45–51. doi: 10.1016/j.cobeha.2020.12.008.

Naselaris, Thomas, Kendrick N. Kay, Shinji Nishimoto, and Jack L. Gallant. 2011. “Encoding and Decoding in FMRI.” NeuroImage 56(2):400–410. doi: 10.1016/j.neuroimage.2010.07.073.

Naselaris, Thomas, Cheryl A. Olman, Dustin E. Stansbury, Kamil Ugurbil, and Jack L. Gallant. 2015. “A Voxel-Wise Encoding Model for Early Visual Areas Decodes Mental Images of Remembered Scenes.” NeuroImage 105:215–28. doi: 10.1016/j.neuroimage.2014.10.018.

Nguyen, Mai, Tamara Vanderwal, and Uri Hasson. 2019. “Shared Understanding of Narratives Is Correlated with Shared Neural Responses.” NeuroImage 184:161–70. doi: 10.1016/j.neuroimage.2018.09.010.

Polyn, S. M. 2005. “Category-Specific Cortical Activity Precedes Retrieval During Memory Search.” Science 310(5756):1963–66. doi: 10.1126/science.1117645.

Richter, Franziska R., Rose A. Cooper, Paul M. Bays, and Jon S. Simons. 2016. “Distinct Neural Mechanisms Underlie the Success, Precision, and Vividness of Episodic Memory.” ELife 5:e18260. doi: 10.7554/eLife.18260.

Rissman, Jesse, and Anthony D. Wagner. 2012. “Distributed Representations in Memory: Insights from Functional Brain Imaging.” Annual Review of Psychology 63(1):101–28. doi: 10.1146/annurev-psych-120710-100344.

Rugg, Michael D., and Kaia L. Vilberg. 2013. “Brain Networks Underlying Episodic Memory Retrieval.” Current Opinion in Neurobiology 23(2):255–60. doi: 10.1016/j.conb.2012.11.005.

Smith, Stephen M., and J. Michael Brady. 1997. “SUSAN—A New Approach to Low Level Image Processing.” 34.

Song, Hayoung, Emily S. Finn, and Monica D. Rosenberg. 2021. “Neural Signatures of Attentional Engagement during Narratives and Its Consequences for Event Memory.” Proceedings of the National Academy of Sciences 118(33):e2021905118. doi: 10.1073/pnas.2021905118.

Sprague, Thomas C., Edward F. Ester, and John T. Serences. 2016. “Restoring Latent Visual Working Memory Representations in Human Cortex.” Neuron 91(3):694–707. doi: 10.1016/j.neuron.2016.07.006.

St-Laurent, Marie, Hervé Abdi, and Bradley R. Buchsbaum. 2015. “Distributed Patterns of Reactivation Predict Vividness of Recollection.” Journal of Cognitive Neuroscience 27(10):2000–2018. doi: 10.1162/jocn_a_00839.

Walther, Alexander, Hamed Nili, Naveed Ejaz, Arjen Alink, Nikolaus Kriegeskorte, and Jörn Diedrichsen. 2016. “Reliability of Dissimilarity Measures for Multi-Voxel Pattern Analysis.” NeuroImage 137:188– 200. doi: 10.1016/j.neuroimage.2015.12.012.

Xiao, Xiaoqian, Qi Dong, Jiahong Gao, Weiwei Men, Russell A. Poldrack, and Gui Xue. 2017. “Transformed Neural Pattern Reinstatement during Episodic Memory Retrieval.” The Journal of Neuroscience 37(11):2986–98. doi: 10.1523/JNEUROSCI.2324-16.2017.

Yu, Qing, and Won Mok Shim. 2017. “Occipital, Parietal, and Frontal Cortices Selectively Maintain Task-Relevant Features of Multi-Feature Objects in Visual Working Memory.” NeuroImage 157:97–107. doi: 10.1016/j.neuroimage.2017.05.055.

Zadbood, A., J. Chen, Y. C. Leong, K. A. Norman, and U. Hasson. 2017. “How We Transmit Memories to Other Brains: Constructing Shared Neural Representations Via Communication.” Cerebral Cortex 27(10):4988–5000. doi: 10.1093/cercor/bhx202.

